# Craniofacial allometry is a rule in evolutionary radiations of placentals

**DOI:** 10.1101/513754

**Authors:** Cardini Andrea

## Abstract

It has been suggested that larger species of mammals tend to become long-faced when they diverge in size during an evolutionary radiation. However, whether this allometric pattern, reminiscent of ontogenetic changes in skull proportions, is indeed a rule has yet to be thoroughly tested. Using ~6000 adult specimens from 14 phylogenetically well separated and ecomorphologically distinctive lineages, 11 orders, and all superorders of the placentals, I tested each group for positive craniofacial allometry (CREA). The evidence supporting CREA is overwhelming, with virtually all analyses showing proportionally longer faces in bigger species. This corroborates previous studies in other groups, consolidates CREA as a pervasive morphological trend in placental evolution and opens important research avenues for connecting micro- and macro-evolution. If found in even more lineages of non-placental mammals, confirmed in birds, and possibly discovered in other tetrapods, CREA could become one of the most general rules of morphological evolution in land vertebrates.

---

Scale has a vast range of consequences on a variety of aspects of biology including body proportions and the shape of specific anatomical regions (1–3). In morphology, allometry refers to size-related variation in ontogeny and evolution (4), and can act both as a constraint of change or a promoter of novelty along lines of least evolutionary resistance (2, 5–7). Mammals offer particularly interesting examples to explore the role of size and allometry in evolution, as they vary in body mass more than any other class of vertebrates: the smallest shrew or bat weighs just a few grams, while the blue whale may exceed 150 tons and is the largest animal of all times. Mammals also tend to have larger brains compared to other tetrapods, and are defined by a large number of derived traits in head anatomy (heterodont dentition, secondary bony palate, complex turbinates, paired occipital condyles etc.). Thus, the origin of craniofacial differences and allometric variation is central to the study of mammalian biology and evolution.

Adults of larger species, in a group of closely related mammals, tend to have relatively longer faces and smaller braincases (5, 8). This macroevolutionary pattern of cranial evolutionary allometry (CREA (9)) was first described as mammalian trend by Radinsky (8) and has later been hypothesized to be a rule of morphological evolution (5, 9). CREA has been mainly explored in placentals (5, 10), but occurs also in marsupials, such as didelphids (8) and kangaroos (9), and more recently, but less conclusively, has been supported even in birds (11–13). However, before it can join the list of established evolutionary ‘rules’, stronger evidence from a larger number of species and lineages must be obtained.

In this context, it is important to bear in mind that the term “rule” is used in a loose sense to refer to broad patterns, that are common in a group of organisms but may vary in magnitude and can have exceptions (14). Despite possible differences and exceptions, however, they represent consistent evolutionary trends across a large number, or even the majority of taxa, in a lineage. Thus, Bergmann’s, Allen’s, Rensch’s and other rules, including CREA, are not strict laws describable by constants of nature (14), but general patterns, whose taxonomically contingent occurrence in a vast number of cases suggests commonalities in the mechanisms behind them. Causal processes are seen by some as a fundamental requirement for proposing a rule (15). Indeed, that may be true in specific historical contexts, such as the original formulation of Bergmann’s rule (15), but does not apply to the vast majority of evolutionary rules, that are best described as *explananda* rather than *explanantia (14)*. In fact, mechanisms tend to be often speculative and based on correlational evidence. Even in the case of Bergmann, possibly the best known evolutionary rule in mammals, its explanation is often presented as a fact, but is largely still debated (16).

That rules are first accurately verified and documented seems more important than proposing explanations before the pervasiveness of a pattern has been robustly assessed. Thus, focusing on placentals, the first and main lineage in which CREA has been described, and using a sample of almost six thousands specimens and 235 species (Table S1), I will examine whether this pattern is supported in 14 phylogenetically well separated and ecomorphologically disparate lineages from 11 different orders, representing all four superorders of placentals. The study follows the traditional allometric school of Huxley and Teissier (4, 17), and employs major axis (MA) regressions and comparative methods on species average measures of facial and braincase overall size, ventral and dorsal lengths. In total, more than 2000 allometric slopes will be estimated to explore the sensitivity of results to: the choice of the morphometric descriptor; the taxonomic level (with both supra- and, whenever enough species are available, infra-generic analyses); sexual dimorphism, if present and large; the inclusion or exclusion of species with very small samples (N≤5); and differences in branch lengths in phylogenetic trees (corresponding to models varying from star radiation and early diversification to intermediate and late diversification, as in (10)).

The anatomical points used to compute (a) facial and braincase centroid size are shown in Figure 1 together with (b) the lengths of the nasals and cranial vault (dorsal view), and (c) those of the palate and cranial base (ventral view). Support for CREA is found when the increase in facial size during an evolutionary radiation is faster than that of the braincase, which is indicated by slopes >1 in MA regressions of facial onto braincase measurements. The main results are summarized in Table S2 and slopes are shown in Figures 2-5. The vast majority of analyses and taxa support the prediction that the face is characterized by positive allometry (slope >1) relative to the braincase. The relationship between facial and braincase measurements is strong with almost 80% of regressions accounting on average for more than 2/3 of variance (R2) in the data. The average of slopes across evolutionary models in each set of analyses (Table S2) is >1 in 96% of cases, >1.2 in 75% and >1.5 in 34%. Some groups have stronger patterns and steeper slopes (e.g., most primates and sometimes the Africant megabats), and others show weaker trends with shallower slopes (e.g., often armadillos or, sometimes, canids and ungulates). However, virtually all lineages, regardless of the type of measure of craniofacial proportions and the inclusion or exclusion of smaller samples, indicate that, among closely related species, the size-related trend of facial elongation and braincase reduction is clear and consistent. Thus, as for Bergmann’s rule (16), the strength of the trend varies from case to case, but the pattern is overwhelmingly supported.

**Fig. 1.**
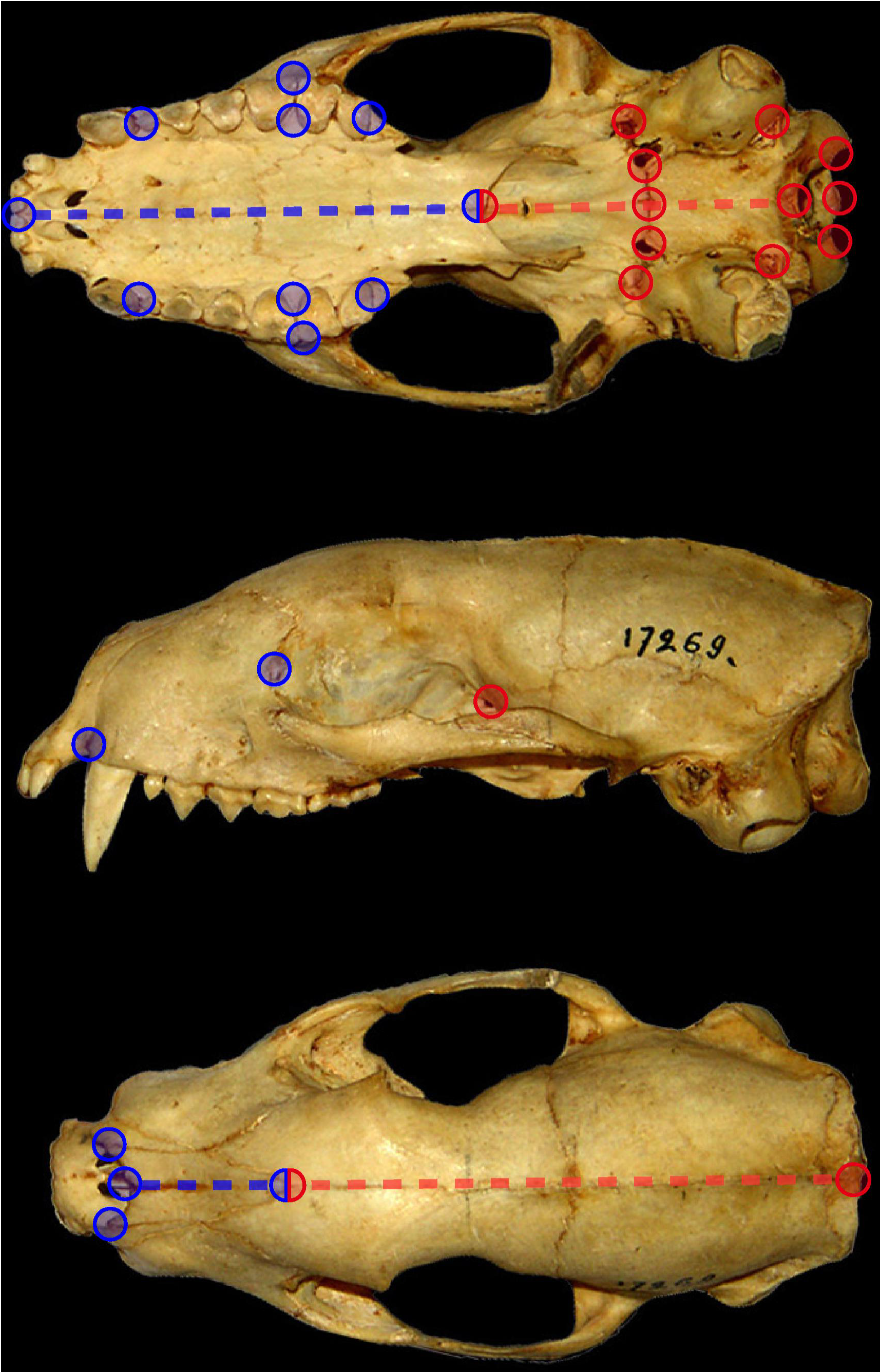
Measures. Anatomical landmarks and linear distances used to measure the size and length of the face and braincase.

**Fig. 2.**
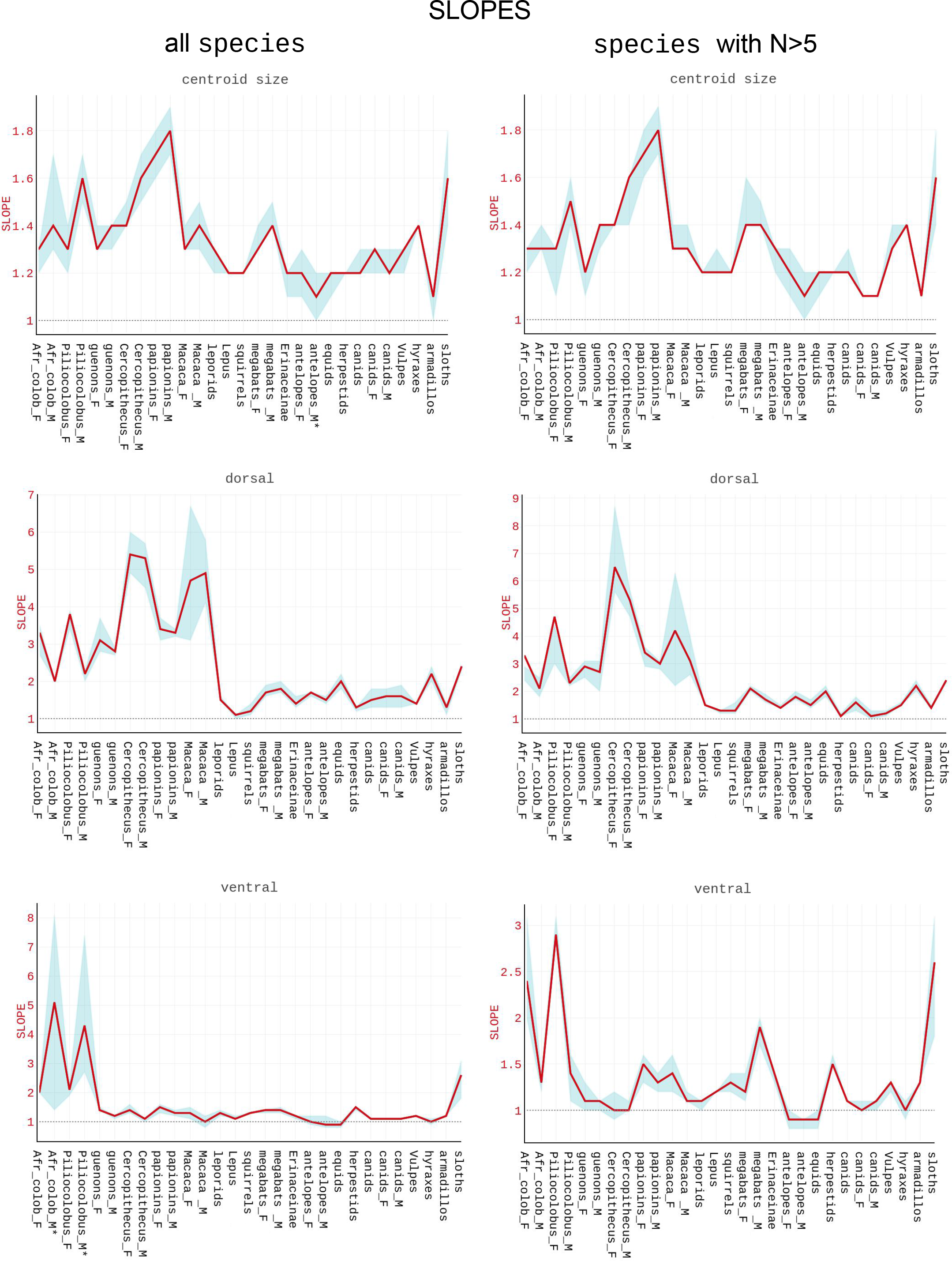
Summary of slopes. Profile plots of average slopes (red lines) using all species or the reduced dataset with larger species samples (N>5); the light blue shaded area shows the range of variation (10th and 90th percentiles) of slopes estimated using regressions with different branch lengths; the horizontal dotted line indicates isometry (slope =1). Abbreviations: Afr_col, African colobines; F, females; M, males.

**Fig. 3.**
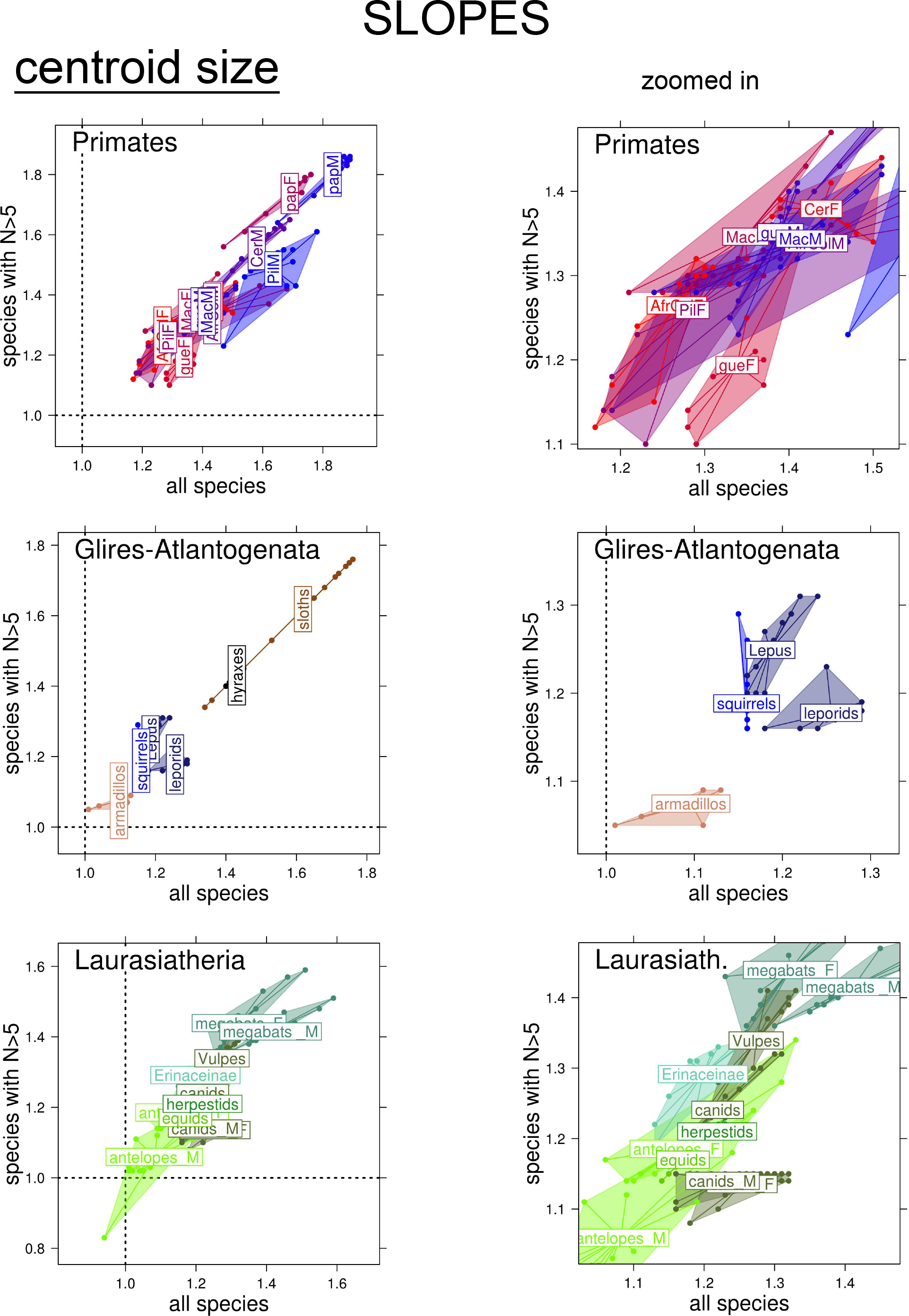
Slopes for centroid size. Scatterplots of slopes using different branch lengths in comparative analyses. Slopes from analyses of the full dataset are on the horizontal axes and those of the reduced dataset (including only species with N>5) are on the vertical axes. Dotted lines indicate isometry. Scatterplots are grouped by taxa. Abbreviations use the first three letters of the genus name; for African colobines, guenons and papionins the abbreviations are respectively Afr_col, gue and pap. F and M indicate respectively females and males in analyses with separate sex.

**Fig. 4.**
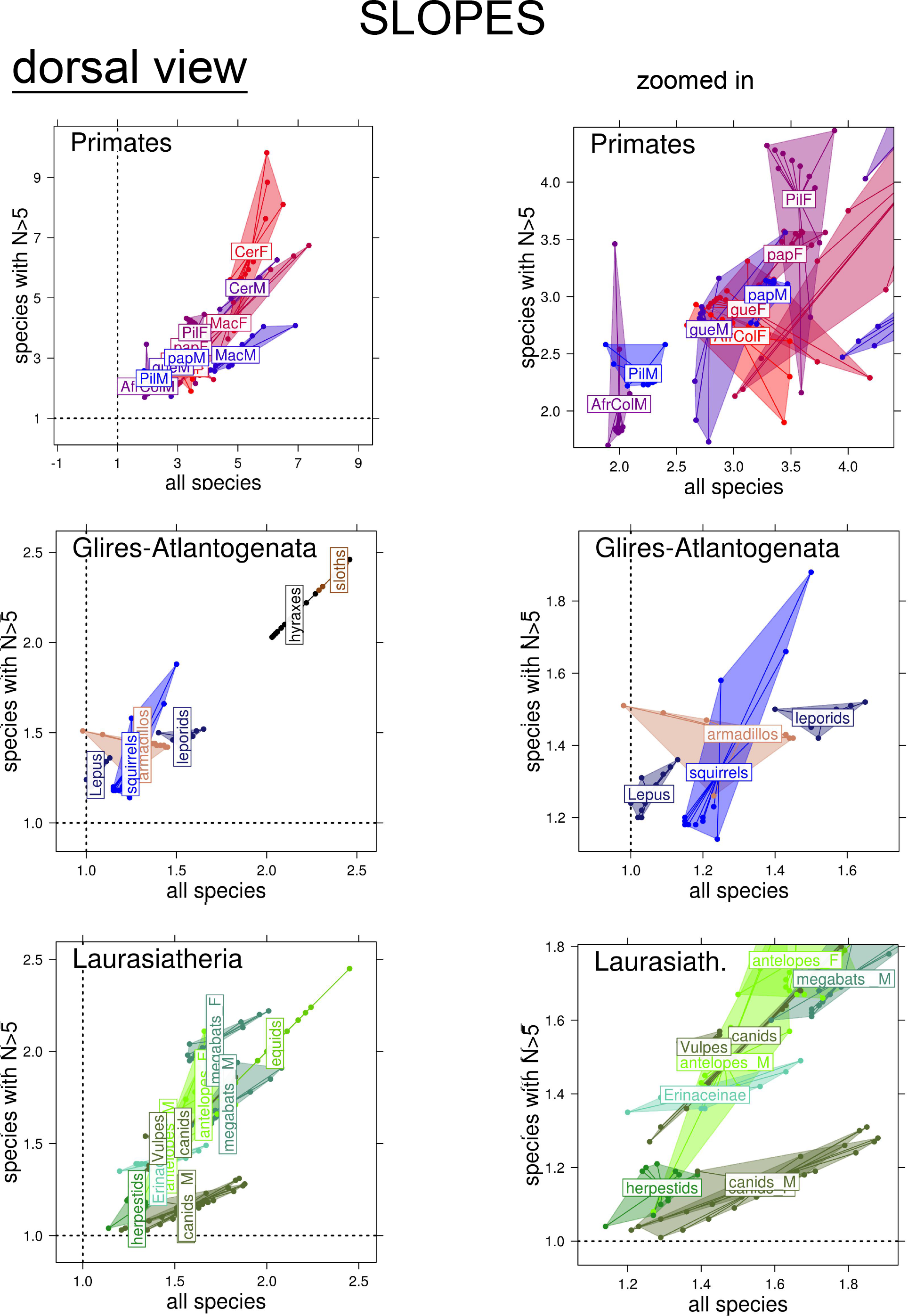
Slopes for dorsal lengths. Scatterplots of slopes based on different datasets, models and taxa. See Fig. 3 for full legends.

**Fig. 5.**
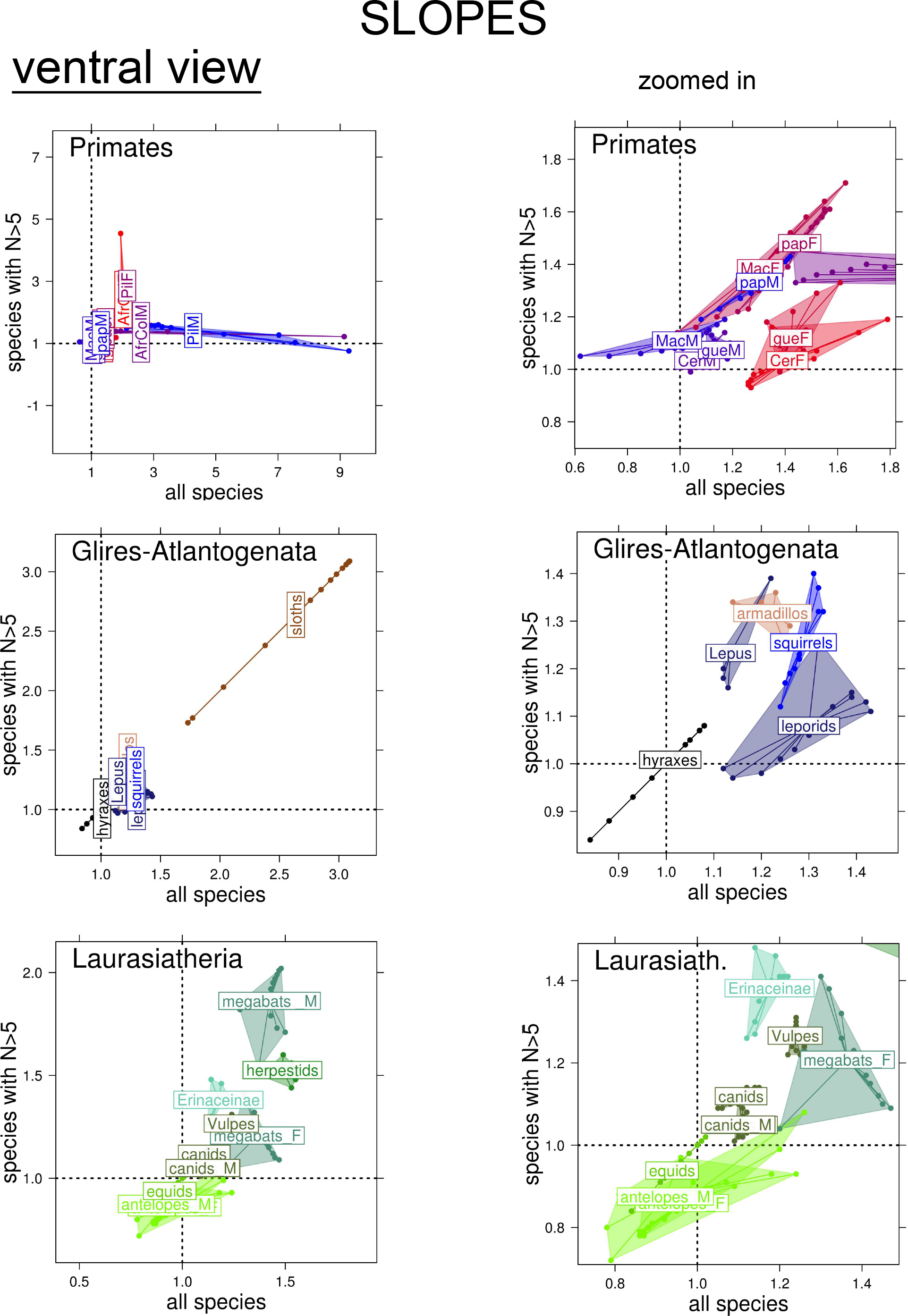
Slopes for ventral lengths. Scatterplots of slopes based on different datasets, models and taxa. See Fig. 3 for full legends.

There are some exceptions to CREA, and they mostly occur in a very limited number of lineages using lengths in ventral view. In ungulates, African antelopes and equids, the palate and cranial base lengths can increase isometrically in bigger species or sometimes even with a slight shortening of the face in relation to the cranial base. Radinsky (8) also analysed equids, as well as the whole bovid family, but did not find them to depart from the general trend of positive allometry. However, he could only use ordinary least square models (that do not take the non-independence due to phylogeny into account), employed a slightly different measure of braincase size, and only analysed the ventral view of the cranium. He also sampled different species, presumably including fossils in the high-crowned equids and using 18 species to represent the whole diversity and disparity of the bovids. His study was based on a single individual for each species and did not take possible sex differences into account. Thus, the difference between my results and Radinsky’s findings only concerns a minority of cases using ventral lengths in two groups, that did not fully overlap across our analyses and may have different issues in terms of sampling error and statistical inaccuracies.

Besides ungulates, and again only using ventral lengths, also canids (as in Radinsky (8)), hyraxes, *Cercopithecus* monkeys and male macaques support CREA less strongly, as positive allometry is on average very moderate in these groups and sometimes isometry is found. However, results that might contradict CREA represent overall less than 10% of the total, with in fact dorsal lengths as well as analyses of centroid size consistently confirming positive allometry of the face in all taxa, including ungulates, hyraxes, canids and all monkeys.

The main conclusion of an overwhelming support for CREA is therefore robust. However, potential exceptions to the rule, such as ventral lengths in ungulates and few other groups, provide interesting examples of how different taxa, and cranial regions, may have departed from the general trend. This, and the variability of slopes, suggests that CREA acts as a widespread constraint (in terms of direction but not necessarily of speed) of changes in craniofacial proportions during evolutionary radiations. The constraint is of moderate strength, and the rule is flexible and can be modified, or even reversed, by natural selection. For instance, in ungulates, the pressure for breaking the rule in ventral cranial proportions could have been the need of housing large hypsodont teeth, whose size might be relatively larger in small herbivores, forcing the palate to be proportionally long despite a modest cranial size. Tamagnini et al. (10) also speculated that teeth might be important in modifying the rule, as the main exception among pantherines was the proportionally long palate of the relatively small clouded leopard, a species with exceptionally long canines and therefore very deep and long dental roots. In human evolution, in contrast, selective pressures for evolving large brains, together with tool use, cooking, and other forms of food processing, made large teeth unnecessary and probably contributed to revert CREA (5). That thus led to the distinctive morphology of modern humans, with their disproportionately large braincase and orthognathic face.

The robust support for CREA in 14 distantly related, and ecomorphologically disparate, placental lineages makes it unlikely that the pattern arose independently so many times. In fact, this analysis, the largest until now in terms of samples and taxonomic groups, together with Radinsky’s (8) pioneering work, and a number of studies reporting similar craniofacial allometric patterns within one or or another specific lineage of mammals, provide evidence for CREA in: four large taxa of primates (8), as well as four families of carnivores (8–10); several groups of rodents (5, 18–20), and the main living family of the lagomorphs, the leporids (8); three lineages of arctiodactyls (5, 8, 21), and at least one of perissiodactyls (8); one of the main lineages of erinaceomorphs (8) and one of bats (5); two families of Afrotheria (the tenrecs (8), besides the hyraxes analysed here); two out three of the main xenarthran clades; and at least two families of marsupials (8, 9). Thus, parsimony suggests that most other placentals might follow CREA, that, if found in even more lineages of marsupials (and maybe also in the echidnas), confirmed in birds (11–13) and possibly discovered in other land vertebrates, could become one of the most general rules of morphological evolution in the tetrapods.

Probably the most important aspect not to forget in future studies of CREA concerns the choice of an appropriate taxonomic level for the analysis. Radinsky (8) largely focused on families. That has the advantage of apparent consistency and might seem more objective. However, neither taxonomy nor evolutionary time can provide on their own a simple criterion to select the level at which CREA should be tested. Some families may be more conservative in cranial form and others show larger disparity. And even closely related lineages may evolve along different morphological trajectories. Cercopithecines and colobines are both members of the family Cercopithecidae, but cercopithecines, and especially papionins, are mostly omnivorous and tend to have on average long snouts, whereas the predominantly folivorous colobines are typically orthognathic. Such large differences in ecology and average cranial proportions between subfamilies or tribes of Old World monkeys make the cercopithecid family as a whole an unlikely target for testing CREA. Tests within its subclades, in contrast, are appropriate and have until now produced support for the rule. Similarily, CREA occurs both within the felids (10) and the viverrids (8), whereas a test performed after pooling the generally larger but short-faced hypercarnivorous cats and the smaller long-faced omnivorous genets would not make much sense despite their close phylogenetic relationship. This is not only because, as the Bermann’s rule (16), CREA is defined as a trend found in closely related species and populations (5). It is also, and more importantly, because pooling lineages that diverged so much in ecology, that their formerly common cranial bauplan becomes different in terms of typical craniofacial proportions, would produce a meaningless test of CREA, clearly outside the realm in which the rule has been proposed and described (5). Indeed, a sharp drop in the fit of a single regression line in a larger lineage, compared to separate regressions within its sub-clades, might often be a clue that differences have become too large for pooling the clades.

In contrast, within a group sharing a common cranial bauplan, CREA might be tested and supported (with more or less strength) or rejected. If CREA is supported, that will be as an average trend in that group, despite possible and potentially interesting exceptions. Among big cats (subfamily Pantherinae), as mentioned, the clouded leopard is relatively long-faced despite being smaller than other species (10), thus deviating from the general trend probably because of its very long canines (proportionally longer than in any other living felid). In the case of simple bivariate regressions, as in this study, the magnitude of the deviation from the main regression line can be taken as an estimate of how unusual a species is for craniofacial proportions compared to its closest relatives, exactly in the same way as one uses the encephalization quotient to infer unusually large (or small) brains relative to body mass (22). Indeed, as the pace of evolution varies across groups and structures, there will not be a simple unique taxonomic level for testing CREA, that fits all taxa. To make the taxonomic level less approximate, one could try to relate the strength of CREA (and its likely refutation above a certain degree of variation) to morphological disparity across groups within a larger clade.

As anticipated, for now, one can only speculate about the processes behind CREA. The resemblance to an almost universal mammalian trend of within species postnatal ontogenetic change (5), in which the rate of facial growth is faster than that of the braincase, making the face of adults proportionally longer, is evident and intriguing. One might assume that, if this similarity is based on homologies, it could provide an evolutionary developmental link between micro- and macro-evolutionary mechanisms of morphological variation. That adults of domestic mammals, selected to be smaller than their wild counterparts, in general originally tended also to be somewhat paedomorphic and short-faced seems consistent with a relationship between growth and the origin of evolutionary change (5). Also, a genetic correlation between craniofacial variation and body mass has been demonstrated in primates and suggested to provide a potential mechanistic explanation for CREA (23). Nevertheless, as we and others discussed in previous papers (5, 8, 18), dietary pressures and biomechanical adaptations might provide alternative and non-mutually exclusive explanations of CREA. In this respect, showing that facial elongation in larger species is adaptive to, for instance, climate (24) does not refute CREA, as CREA is purely about the pattern and does not assume any specific mechanism. It does not even rule out an hypothetical explanation in terms of genetic and developmental architecture (23, 25), which might on one hand constrain to a larger or smaller degree evolvability (26), but, within this narrower range of potential directions of variation, can also provide a simple ‘evolutionary trick’ to speed up change and produce adaptations along an allometric line of least evolutionary resistance (6, 7).

Evolution tinkers with available mechanisms and often exploits already existing developmental processes to produce morphological variation (27). Both brain and metabolism scale negatively with body mass in mammals, thus making negative allometry of the braincase somewhat expected (5). However, ontogenetic trajectories usually diverge progressively across species as they radiate, as shown in a variety of mammals from rodents (28) to hominins (29). Therefore, interspecific allometries may share aspects of a common ontogenetic pattern but are unlikely to be produced by simple extension or truncation of size-related developmental trajectories. To make the picture more complicated, epigenetic factors, such as hormone levels, muscle development and dietary change, act upon bone shape during growth, adding a layer of plasticity to the phenotypic output of regulatory networks and the potential constraints set by gene linkage and pleiotropy (5, 23). Discovering the process (or the multiplicity of processes) behind evolutionary rules is clearly a challenge, but lacking an explanation does not diminish the importance of robustly documenting patterns (16). In this respect, CREA is now a big step closer to becoming a well supported evolutionary rule of morphological change in placentals and probably other groups of mammals and vertebrates.

**Table 1.**
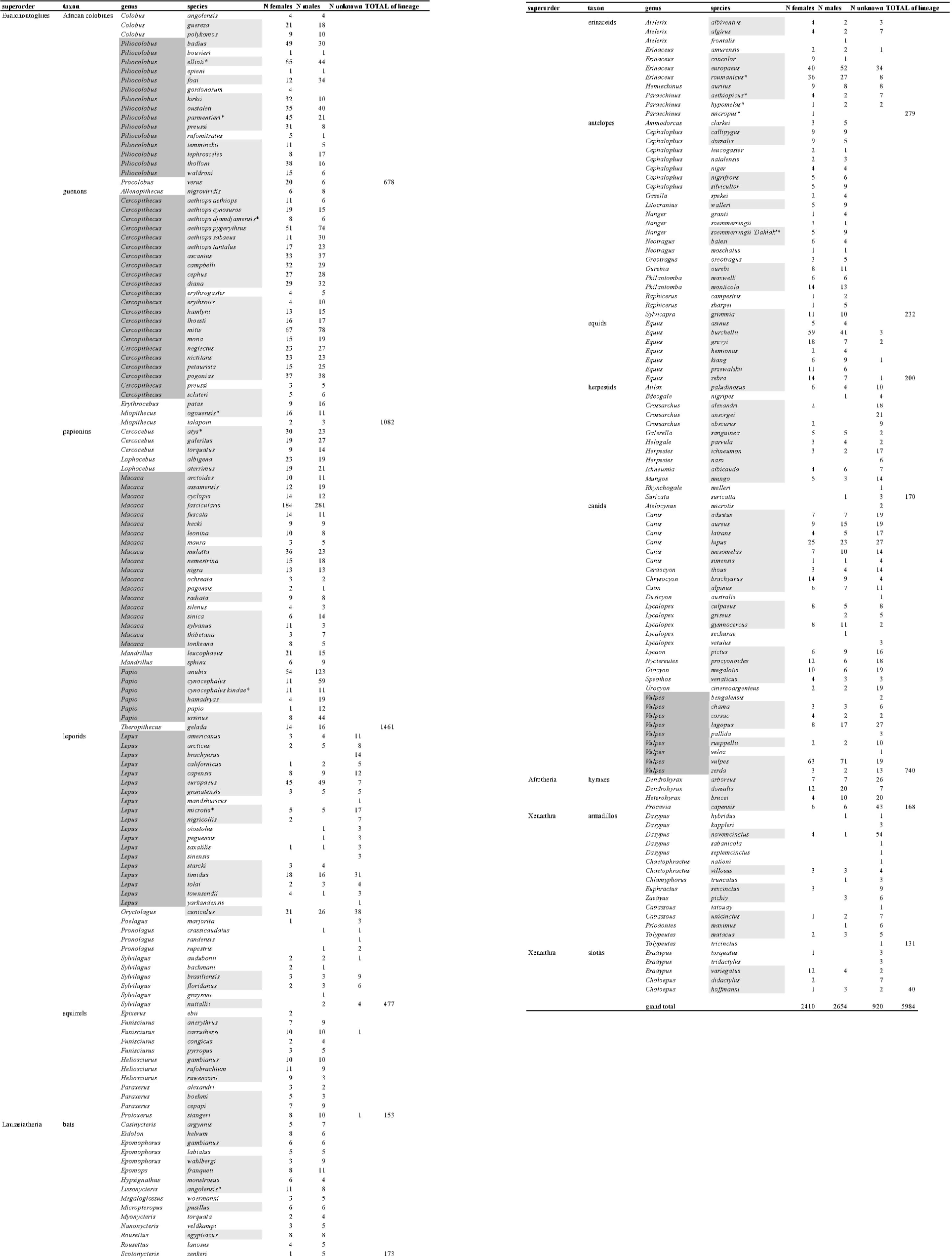
Sample Composition. A dark background is used for genera with at least nine species, which were tested on their own as well as within the main lineage to which they belong; a light grey background emphasizes species with N>5 (at least in one sex, if analysesd using seperate sexes); * marks species not available in the phylogeny and therefore excluded from comparative analyses.

**Table 2.**
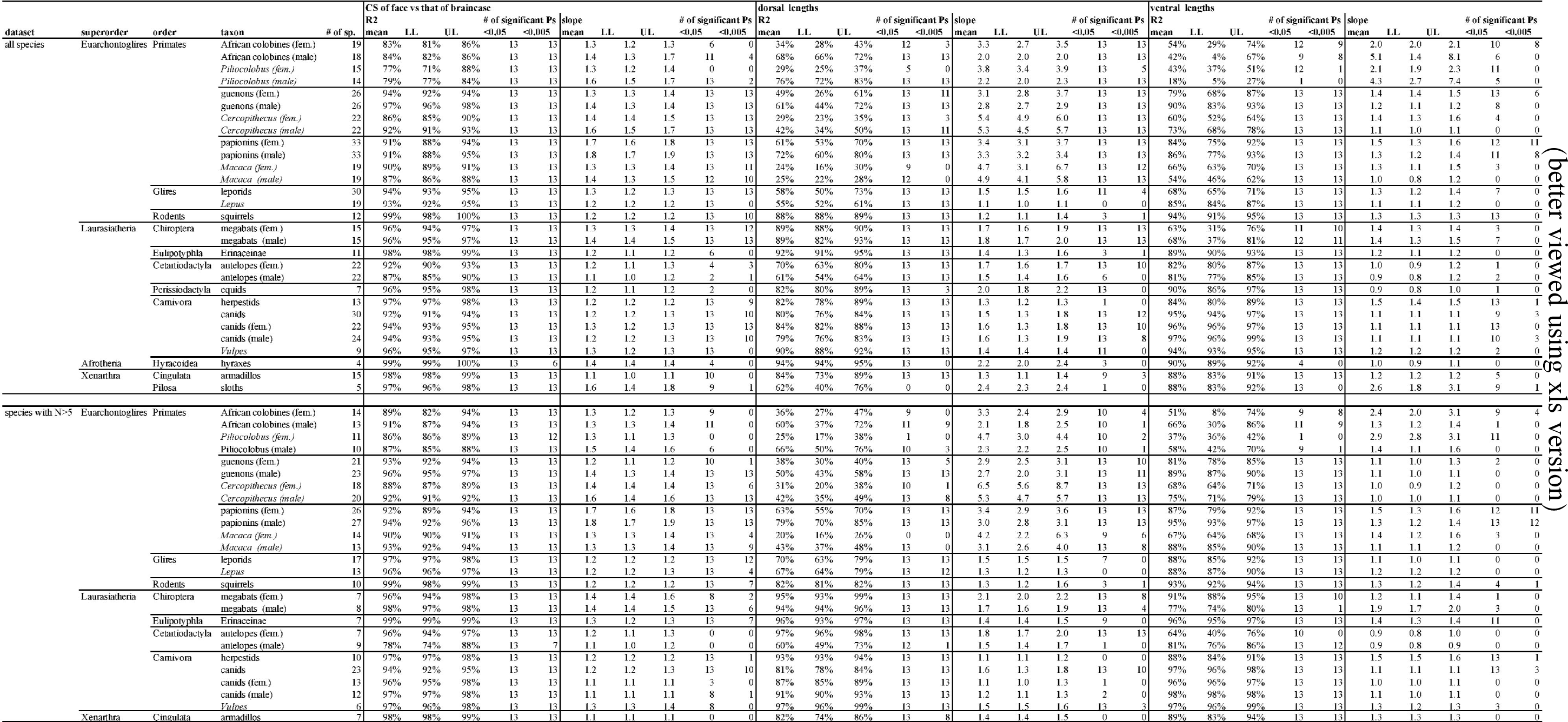
MA regressions. Summary or results (for each t.axon. 13 regressions using differem branch lengths in comparative analyses): number (#) of species in a taxon; rn an of regression estimates for R2 and slope together with their respective 10th and 90th perce1uiles (LL aOO UL); numberof significam tests (R2 or slope different to I) using P<0.05 and P<0.05 as significance thresho

## Acknowledgements

New data, which are published for the first time in this paper, were collected at Museo Civico di Storia Naturale di Milano, Natural History Museum Vienna, National Museum Prague, Museum für Naturkunde, Museum National d’Histoire Naturelle, Hungarian Natural History Museum, Naturhistoriskariksmuseet. I owe a huge thank to all curators, collection managers and staff, including the SYNTHESYS administrators, whose help and support was fundamental. I am also deeply grateful to a number of colleagues who provided technical help and advice. With anticipated apologies for certainly forgetting to explicitly mention several of the many people whom I am in debt with, as well as for listing names in quasi-random order, I would like to thank a lot: Wim Wendelin, Emmanuel Gilissen, Giorgio Bardelli, Görföl Tamás, Gabor Csorba, Virginie Bouetel, Aurelie Verguin, Frank Zachos, Alex Bibl, Krapf Andrea, Petr Benda, Karel Kaderàbek, František Vacek, Manja Voss, Christiane Funk, Detlef Willborn, Mayer Frieder, Daniela Kaltoff, Irene Bisang, Emily Dock Åkerman, Jessica Joganic, Aurélie Siberchicot, David Katz, Mike Collyer, David Warton, Sara Taskinen, Paul O’Higgins, Sarah Elton, David Polly, Krish Seetah, Dan Franklin, Carlo Meloro, Diego Fontaneto and Alessandro Minelli. A special thank also to the greatly missed Colin Groves, for his fundamental contribution to interpreting the preliminary analyses of wild asses, and to Anderson Feijó, who helped with the identification of some of the armadillos. My gratitude goes also to the American Society of Mammalogists, for inviting me to present the preliminary results of this study at their annual conference and for covering all the costs of my trip to the US. This paper is dedicated to the memory of Mario Zambarbieri and Paolo Tongiorgi, my greatest mentors and friends, as well as to that of Colin Groves, a most extraordinary mammalogist, and of Leonard Radinsky, first discoverer of CREA, whose important role I would have acknowledged well before, had I found earlier his surprisingly little cited and yet fundamental contribution on patterns in mammalian morphological evolution.

## Funding, author contributions, competing interests, and data

Data collection was funded by SYTHESYS (BE-TAF-2861, SE-TAF-4409, AT-TAF-4816, CZ-TAF-4817, HU-TAF-4818, DE-TAF-4819, FR-TAF-4820) for all lineages except most primates, that were measured for a previous project with Sarah Elton, funded by the Leverhulme Trust (F/00128/T). This study was designed, all data were collected, analyses done, and the paper written by A. Cardini. I declare no competing interests or conflict of interest. Data used for all analyses are available from the author upon request.

## Materials and Methods

### Samples, classification and phylogeny

5984 adult specimens from 14 lineages belonging to 11 orders, representative of the four placental superorders, and a total of 235 species are studied using specimens from museum collections. Most individuals come from wild populations. Zoo specimens are used only when wild ones are absent or rare in collections, and exclusively if they do not show unusual morphology. Antelopes, bats and primates are analysed using separate sexes because most species show large sex differences (5, 30, 31). In canids, the degree of sexual dimorphism varies depending on the species and therefore analyses are done using both pooled samples, as well as separate sexes. All other lineages, with small and generally negligible sex differences in most species, are analysed regardless of sex. This is done also for *Vulpes*, because information on sex was missing in too many of its species for separate sex analyses. Thirteen species (5.5% of the total) are excluded from comparative tests, because they are not available in the phylogenetic tree.

Allometry is tested at or below family level, focusing, whenever possible, on species rich taxa with a relatively conserved cranial morphology (common “cranial bauplän” *sensu* Cardini & Polly (5)). Analyses are replicated also at generic level (e.g., in *Lepus*, *Macaca*, *Vulpes* etc.), when a relatively large number of species (nine or more in the full dataset) is available in a genus. Sampling is less extensive in Xenarthrans, that are not well represented in the European museums, where most of the data were collected. For this reason, both armadillos and sloths are analysed at respectively ordinal (Cingulata) and subordinal (Folivora) levels, and confirmatory studies within more homogeneous subgroups, such as family, subfamily or tribe, are desirable especially in the more diverse and older armadillo lineage. Overall, 29 taxa (including sub-groups) are analysed in the full dataset and all of them but three (in which all species had large samples) are re-analysed using only species with larger samples (see below).

Table S1 describes the sample composition. Museum provenance for antelopes, African fruit bats and African tree squirrels, mongooses and primates is detailed in previous studies (5, 31–33).

New data (see Acknowledgements for the lists of museums) were collected for leporids, hedgehogs, equids, canids, hyraxes, armadillos and sloths, and also for meerkats and Dahlak gazzelles, which were added to the already published samples of, respectively, herpestids and antelopes. The classification follows Wilson & Reader (34), which is adopted in most museums^1^. The main exceptions are: primates, classified according to Grubb et al. (37) (updated for red colobus, subgenus *Piliocolobus*, following Ting (38)); vervet monkeys, with separate subspecies following the 10Ktree (39); and *Papio cynocephalus kindae* and Dahlak gazelles (an island population of *Nanger soemmeringii (40)*), that are kept separate from their parental species, because they represent highly distinctive (40, 41) dwarf populations (37, 40, 41).

Comparative analyses employ the chronogram of Bininda-Emmonds et al. (42). However, for for primates, canids and equids, the more recent and better resolved consensus chronogram of the 10KTrees phylogeny is used (39).

### Measurements

3D coordinates of cranial landmarks (Fig. 1) are obtained using a 3D digitizer, as detailed in Cardini & Polly (5). Compared to their anatomical configuration, five landmarks (anterior and posterior tips of acoustic meatus, tip of paraoccipital process, infraorbital foramen, notch of postorbital process) are excluded, as they are not available in all taxa; another landmark, zygo-temp inferior, is removed in order to de-emphasize facial width and focus mostly on length. Facial and braincase configurations are relatively independent, as they only share two boundary landmarks (palatine and nasion). As shown in previous work (5, 32), repeatability is high, and especially so for size measurements in macroevolutionary analyses using species averages.

To test CREA, the species average sizes of the face and the braincase (Fig. 1) are estimated using three paired sets of measurements calculated from the 3D landmark coordinates: 1) centroid size (CS - i.e., the square root of the sum of squared distances of each landmark from the centroid of that configuration of landmarks) of the facial landmarks *versus* CS of the braincase landmarks; 2) the lengths of the nasals (rhinion to nasion interlandmark distance - ILD) relative to that of the cranial vault (nasion to inion ILD); 3) the length of the palate (prosthion to palatine ILD) relative to that of the basicranium (palatine to basion ILD). For brevity, the three pairs of measurements are called overall size (1), and dorsal (2) and ventral (3) lengths of the face and the braincase. As typically larger species have larger crania (5, 9, 10), CREA is framed simply as a size-related trend of craniofacial variation, without specifically stressing that the focus is on cranial dimensions. This convention will be adopted also in this study, although more rigorously one should say that the face increases its size faster than the braincase during evolutionary radiations, so that, among closely related species, those with larger crania tend to have proportionally bigger and longer snouts.

Allometric regressions are performed using all species but also repeated including only species with sample size N>5. Although sampling error is less of a concern in macroevolutionary analyses of large interspecific differences (8, 9), this ‘reduced sample’ of approximately 70% of all species allows to explore the sensitivity of results to the inclusion of species samples of just one or very few individuals (10, 43).

### Statistical analyses

Allometry is tested with a major axis regression (MA) of log-transformed data, following the traditional school of Huxley (4) and the guidelines of Warton et al. (44). This approach has already been used in one of the three sets of analyses of Tamagnini et al. (10), and has the advantage of narrowly focusing on the main aspects under consideration, the relative size of the face and braincase, leaving out taxon-specific features of craniofacial morphology that are irrelevant to CREA but may be captured by more complex geometric morphometric analyses.

To take non-independence due to phylogeny into account, phylogenetic independent contrasts (PICs - (45)) were used in MA regressions. As in previous studies (5, 10), a Brownian motion (BM) model of evolutionary change is employed (45). No search for an optimal evolutionary model is done, because, when a group contains relatively few species, estimates are inaccurate (46, 47) and results based on BM are robust to model selection (48). However, to explore if results of comparative analyses specific to this dataset are truly relatively insensitive to the choice of the evolutionary scenario, I replicate MA regressions using many different branch lengths (10).

These range from extremes, such as a star radiation (i.e., an ordinary non-comparative least square regression, including also the species not available in the phylogeny) as well as a tree with all branch lengths equal to one (punctuated equilibrium model), to more or less pronounced variations of the original chronogram modified using Grafen’s rho (49). Thus, by increasing this parameter from an initial value close to zero, a set of 10 trees is build that vary from a tree corresponding to an early radiation model (quasi-star radiation) to one with just a few long branches implying recent radiations in few main subclades (10). For the non-comparative ordinary MA regressions, I employ the robust method of Warton et al. (50). However, in all sets of data, results of ordinary MA regressions (not shown) are almost unchanged regardless of using the robust method or not, which suggests that this option, not available in analyses using PICs, does not make an appreciable difference in this specific study.

MA regressions are used to estimate R2 and slopes. R2 is the percentage of variance accounted for by the relationships between the variables, and it is used to assess the strength of the covariation between face and braincase size. The regression slope estimates the relative change of the face (y axis) and braincase (x axis), and should be >1 if CREA is supported. Although significance is estimated in all regressions for both the magnitude of R2 and equality of slope to one (i.e., the null hypothesis of isometry), it has a limited interest in the context of this study. P values are unlikely to be reliable in a study where power is low in lineages with few taxa (e.g., the equids with just seven living species), and type I errors are inflated by multiple testing. More importantly, and regardless of potential issues with over-interpretation of null hypothesis significance testing (51), the main aim here is to assess the occurrence of a pattern across a large number of taxonomic groups in relation to sampling and phylogeny. Thus, R2 and slopes are summarized for each block of 13 regressions (ordinary least square and comparative analyses using BM, punctuated model or the 10 models based on Grafen’s rho) with averages and trimmed ranges. Trimmed ranges are the lower 10th (lower limit or LL) and upper 90th (upper limit or UL) percentiles of observed R2 and slopes, which are less affected by outliers compared to the minimum and maximum of a set of estimates.

Analyses are conducted in R (52) using the following packages (50, 53–58): adegraphics and plotly (plots), ape and geiger (comparative methods), geomorph and Morpho (centroid size), and smatr (MA regression). The script and dataset used for the main analyses are included in the Supplementary Information.

## Supplementary Text

### RESULTS IN DETAILS

#### Sensitivity to species with small N

Table S2 (Supplementary Information) provides the summary of the main results of MA regressions of the three sets of measurements. The table also reports how many times, out of 13 regressions in a lineage, tests are significant at 0.05 (significant) or 0.005 (highly significant) levels. As discussed in the methods, significance testing has a limited value in this context. Figure 2 shows the most important summary results from Table S2, which are the average, LL and UL of regression slopes.

Similarities in profile plots (Fig. 2) and high correlations between results using all species or just those with larger samples show that both average R2s and slopes are highly congruent regardless of the inclusion of species with smaller samples. More precisely, excluding three cases (see below), correlations for R2s (full dataset *versus* reduced one) range between 0.77 (ventral lengths) and 0.90-0.93 (CS and dorsal lengths respectively); the corresponding correlations of average slopes vary from 0.91 (ventral lengths) to 0.93-0.94 (CS and dorsal lengths respectively).

The three sets of analyses, which, unlike all others, are not in good agreement, when replicated in larger samples only, are males of antelopes, African colobines and red colobus. Assuming that analyses based on only larger samples are more accurate, regressions seem to underestimate CS R2 using all species in male antelopes, while the opposite (i.e., an overestimate) happens for ventral length slopes in male African colobines and red colobus. However, these three cases of incongruence of analyses using the full sample and those based on the reduced dataset represent less than 2% of all analyses. Thus, they do not change the conclusion about the robustness of results regardless of the inclusion of species with smaller samples. This is also confirmed by the scatterplots of slopes (Figs 2-4), in which most of the values are close to the diagonal, as expected if using all species (horizontal axis) or including only largest samples (vertical axis) produce similar estimates.

#### R2

The covariation between facial and braincase length is very strong with average R2s ranging (across all groups and sets of analyses, using either all species or the reduced dataset) from 66% (dorsal lengths) to more than 90% (CS) of variance explained. The relationship is particularly strong using CS, as all regressions produce R2s accounting for more than ¾ of variance and virtually all of them have P<0.005. Using lengths in ventral and dorsal view, R2s explain on average about 2/3 to ¾ of variance, with the lowest R2s being around 20%. However, despite a sometimes weaker relationships between facial and braincase lengths, R2 LLs are larger than 67% (i.e., 2/3 of variance) in most analyses in ventral view (ca. 70% of times) and in about half (45%) of those in dorsal view. The vast majority of the regressions are either significant (P<0.05 in more than 90% of them) or highly significant (P<0.005 in more than 70% of cases). Overall, the strength of the allometric relationship is large or very large in virtually all study groups regardless of sample size, as well as the type of measures used to describe the relative size of the face and braincase.

#### Slopes

Figures 3-5 show slopes of different lineages using convex hulls to better visualize the full range of results in all analyses and taxonomic groups. In these figures, primates are shown on their own, as they contained the largest number of lineages among all orders in the analysis. Primates and Glires belong to the superorder Euarchontoglires but, to avoid adding more results in the already crowded primate scatterplots, representatives of the Glires are shown together with those of the Atlantogenata. Finally, slopes for the numerous lineages used to assess CREA in the Laurasiatheria make up the third set of scatterplots. In all these scatterplots, slopes of the full set of species are on the horizontal axis and those of the reduced dataset (species N>5) are on the vertical axis. The lines within each convex hulls connect the observed slopes in the 13 MA regressions of a given taxon with their average, which is the same reported in Table S2 and Figure 2. In this ‘space-of-slopes’, support for CREA is in the region of values >1, which is the area above and to the right of the isometric lines (dotted lines indicating slope of one). If isometric lines are not shown in a scatterplot, it is because it only includes values >1. This generally happens for scatterplots in the left column, which are the same as in the right column, but they zoom in the region where most taxa lie. In a few groups, such as, for examples equids and hyraxes, slopes appear to lie perfectly on the diagonal of the scatterplot, but this is simply because all species have N>5 and therefore values of slopes reported on the horizontal and vertical axis are the same.

In <u>primates</u>, CS slopes (Fig. 1) are always >1, with the largest values (close to 2) in the papionins and the lowest (ca. 1.2-1.3) among females of African colobines and guenons. Macaques and males of guenons and *Piliocolobus* generally had intermediate slopes (ca. 1.4-1.6). Also using dorsal lengths (Fig. 2), slopes are all >1 in primates, although in this view of the cranium, it is *Cercopithecus* monkeys that tend to have larger slopes (ca. equal to 5-6) and males of African colobines and *Piliocolobus* that mostly have lower values (on average ca. 2-2.5). However, papionins and macaques have large slopes as well (ca. 3-5, on average). Because the smallest slopes estimated using dorsal lengths are generally larger than the largest ones estimated with CS, it seems that CREA is well supported by both types of craniofacial measurements, even if stronger in dorsal view.

Using ventral lengths (Fig. 3), the pattern in primates is more complex, as in a minority of instances (mainly, some of the regressions in male macaques, as well as in females of *Cercopithecus*) slopes are <1. There are also two cases of clearly incongruent results between analyses using all species or only species with N>5. As already discussed, these are males of both African colobines and *Piliocolobus*, for which slopes likely tend to be overestimated when smaller samples are included. Excluding these potential overestimates, the largest slopes (ca. 2-3) are those of the African colobines, regardless of sex and whether all species or just red colobus are analysed. The smallest slopes (slightly >1 or even <1) are found in male macaques, guenons and *Cercopithecus*. However, also in ventral view the vast majority of slopes are >1 and within groups, on average, about as large as using CS. Thus, overall, results from CS, dorsal and ventral views largely support CREA in primates in the vast majority of cases.

Among the representatives of the <u>Glires</u> (African tree squirrels and leporids) and <u>Antlantogenata</u> (armadillos, sloths and hyraxes), slopes are virtually always >1. Using both CS and dorsal lengths (Figs 2-4), sloths and hyraxes have the largest slopes (ca. between 1.5 and 2.5 on average). For CS, armadillos have the smallest slopes (ca. 1-1.1) with representatives of the Glires having slightly larger values (ca. 1.2-1.3 but always less than 1.4). For dorsal lengths, *Lepus* and armadillos have some of the smallest slopes in absolute terms, although virtually always >1. As with primates, the ventral lengths (Fig. 5) show some differences in the pattern of slopes. In Figure 5, sloths still have the largest ones (ca. 1.5-3) and hyraxes the smallest, with an average of one and negative allometry (<1) in a few regressions. In contrast, in the same figure, ventral slopes for the Glires are on average similar to those of CS and dorsal measurements, thus modestly >1 (1.2-1.3 on average). Overall, the vast majority of analyses in the representatives of the Glires and the Atlantogenata produce slope >1 and, as in primates, support CREA using all types of measures.

The <u>Laurasiatheria</u> is the most disparate group of representatives of placental radiations in this study, and indeed includes the largest (equids and antelopes) and smallest (African fruit bats) animals in the sample. Results are again mostly congruent regardless of using all species or just those with N>5. Possibly the main, small, exception is the separate sex analyses of canids using dorsal lengths, that produce smaller slopes in the reduced dataset with species N>5. In general, slopes in the Laurasiatheria vary over a smaller range (from slightly <1 to ca. 2-2.5) compared to the Euarchontoglires and Atlantogenata. Across measurements, the most evident congruence is fruit bats showing the largest slopes (ca. 1.4 to 1.7, on average) and antelopes and equids having some of the smallest ones using CS and especially using ventral measurements (ca. 0.9-1.2 on average).

Thus, with the exception of dorsal lengths, which in antelopes and equids have some of the largest slopes (on average, ca. 1.5-2), CREA is most strongly supported in the lineage with the smallest animals (African fruit bats), and much more weakly in those with the largest ones (antelopes and equids). Indeed, in ventral view, antelopes and equids are the main exception to CREA, with most regressions producing slopes <1 (ranging from slightly less than 0.8 to slightly more than 1).

However, in all other cases (CS and dorsal lengths), even in antelopes and equids, slopes are >1 in almost all analyses, and not only in fruit bats slopes are large, but also in all other taxa (i.e., hedgehogs and all representatives of the carnivores, regardless of the type of measurements) there is a consistent support for CREA, with average slopes varying from ca. 1.2 to 1.5.

That CREA is the dominant pattern in all the lineages in this study is visually evident in Figures 2-5. Indeed (Table S2), CS and dorsal measurements virtually always produce slopes >1, with more than 90% of LL being >1.1. For these two types of measurements, average slopes range overall between slightly less than 1.1 (both CS and dorsal) to 1.8 (CS) and >5 (dorsal), with an average of 1.3 and 2.5 for respectively CS and dorsal lengths across all taxa and sets of analyses. The ventral view also produce a large majority of average slopes >1 (almost 90% of the times, with 75% of average slopes >1.1). However, about 10% of average slopes and 16% of LL were <1, with a few cases (4% - mainly the already mentioned ventral lengths of antelopes and equids) with even UL being <1. Nevertheless, even in this minority of cases suggesting the opposite of CREA, with the face becoming proportionally smaller in the bigger species, average slopes were at least 0.9 and thus close to isometry. Indeed, average slopes using ventral lengths ranged between 0.9 and 3-5 (respectively using all species or only those with larger samples) and in almost all taxa supported CREA, showing that also the palate typically becomes proportionally longer than the basicranium as cranial size increases across closely related species. The overwhelming conclusion is therefore that there is a consistent pattern of craniofacial covariation and the pattern is highly consistent with CREA in virtually all groups, regardless of the type of measurement, sex differences, taxonomic level, inclusion of smaller samples and differences in evolutionary models.

Finally, in terms of slopes significantly different from 1 (almost always >1), CS and dorsal lengths are significant (P<0.05) in about 75% of all regressions, with almost 50% of them being highly significant (P<0.005). In ventral view, however, only 40% of regressions are significant and just 10% reach the high significance threshold. Regardless of the type of measurement used to capture the relative proportions of the face and braincase, in all datasets, the number of significant tests of slopes correlates with the number of species in the lineages, which confirms that lack of significance may be often related to low power in taxa with fewer species (either because they are not species rich - as the living equids, hyraxes etc. - or because of limited taxonomic sampling).

### FUTURE EVIDENCE FROM OTHER GROUPS

Although this study has provided strong evidence in support of CREA as a morphological rule in the evolution of placentals, many other lineages should be investigated to explore this common pattern and its exceptions. Besides expanding the study of CREA to other land vertebrates (more sauropsids, but also amphibians) and exploring monotremes as well as marsupials other than kangaroos (9) and didelphids (8), there is a number of placental radiations that might offer particularly interesting clues on differences and potential exceptions to the rule.

Hominins have already been mentioned and, at least in the lineage that led to humans, they almost certainly represent a strong exception. Pangolins and other toothless mammals might be interesting to better understand if teeth, or their lack, exert a strong influence on CREA (Tamagnini, pers. comm.). For the same reasons, as well as as an example of highly autoapomorphic dietary specialization and evolution of extremely large size, whales are clearly another important group to study, and, although smaller and provided with teeth, dolphins too could be interesting, given their highly derived cranial shape and long snout. Among marine mammals, phocid seals are both speciose and disparate for size, and thus another good target for testing CREA.

Lineages representing the smallest (many rodents, elephant shrews and shrews, echolocating bats etc.) and largest terrestrial species (elephants and their extinct relatives, rhinos etc.) are also good candidates to keep probing the generality of the rule at opposite extremes of size, as well as in relation to highly distinctive morphologies and specialized ecologies. Indeed, one of the few remaining terrestrial giants, the hyppo seems to have recently joined the list of taxa following CREA (21). The study of this lineages employed geometric morphometrics and did not specifically tested for CREA, but the pattern emerged with strength in the analyses. This study also demonstrated the importance of including fossils, as the sample consisted of the only two living species plus the recently extinct Madagascan hippopotamus. Fossils may indeed be crucial to increase taxonomic sampling in species poor taxa and may contribute to reconstruct the history of how CREA evolved, as size diverged during evolutionary radiations.

## References

See main list above.

**Table S1. (separate file: tabS1.xls)**

**Sample composition.**

A dark grey background is used for genera with at least nine species, which were tested on their own as well as within the main lineage to which they belong; a light grey background emphasizes species with N>5 (at least in one sex, if analysed using separate sexes); * marks species not available in the phylogeny and therefore excluded from comparative analyses.

**Table S2. (separate file: tabS2.xls)**

**MA regressions.**

Summary of results (for each taxon, 13 regressions using different branch lengths in comparative analyses): number (#) of species in a taxon; mean of regression estimates of R2 and slope together with their respective 10th and 90th percentiles (LL and UL); number of significant tests (R2 or slope different to 1) using P<0.05 and P<0.05 as significance thresholds.

## Footnotes

1 Recently, Burgin et al. (35) proposed an updated classification, which increases the total number of mammalian species of almost 1000 species. About two thirds of the ‘new species’ are previously recognized taxa elevated to species level. If adopted by mammalogists and museums, the new classification will require an extensive reassessment of taxa in research and collections. However, for the present study, it seems likely that splitting into more species might in fact increase the support for CREA by improving taxonomic resolution among closely related populations, whose frequent size divergence is typically accompanied by a CREA pattern. For instance, in both vervets and red colobus, populations of smaller size tend to be short faced compared to populations with bigger animals (33, 36).

